# Mechanistic Insights into Sphingomyelin Nanoemulsions as Drug Delivery Systems for Non-Small Cell Lung Cancer Therapy

**DOI:** 10.1101/2025.01.15.633158

**Authors:** Emma Ramos Docampo, Jenifer García-Fernández, Inés Mármol, Irene Morín-Jiménez, María Iglesias Baleato, María de la Fuente Freire

## Abstract

Sphingomyelin Nanoemulsions (SNs) are promising drug delivery systems with potential for treating challenging tumors, including Non-Small Cell Lung Cancer (NSCLC), which has poor prognosis and a 5-year survival rate below 5%. Understanding the toxicity mechanisms and intracellular behavior of SNs is crucial for optimizing their therapeutic application. This study aims to investigate the interaction between SNs and A549 lung adenocarcinoma cells, focusing on their cytotoxic effects and mechanisms of cellular toxicity. SNs were synthesized and characterized for size, surface charge, and stability. A549 cells were treated with varying concentrations of SNs, and cellular uptake pathways were assessed using inhibitors of energy-dependent processes. Cytotoxicity was evaluated through an MTT assay to determine the IC50 value after 24 hours. Mechanisms of toxicity, including lysosomal and mitochondrial involvement, were examined using co-localization studies, mitochondrial membrane potential assays, and markers of apoptosis. SNs exhibited rapid cellular uptake via energy-dependent pathways. The IC50 concentration for A549 cells was 0.89 ± 0.15 mg/mL, suggesting favorable cytocompatibility compared to other nanocarriers. At IC50, SNs induced apoptosis characterized by lysosomal damage, mitochondrial membrane permeabilization, and release of apoptotic factors. These effects disrupted autophagic flux and contributed to cell death, demonstrating potential for overcoming drug resistance. Resveratrol-loaded SNs showed enhanced cytotoxicity, supporting their application as targeted drug delivery vehicles. This study highlights the potential of SNs as efficient drug delivery systems for NSCLC therapy, offering insights into their cellular interactions and toxicity mechanisms. These findings pave the way for the rational design of SN-based therapeutic platforms for cancer and other mitochondria-related diseases.

## 1. Introduction

The latest advances in nanomedicine have opened doors to a new era in developing innovative therapies. Using nanocarriers results in more efficient in-tumor drug delivery, thus leading to a higher antitumor effect and a reduction in healthy tissue damage [1–3]. Among the great variety of materials for the development of nanocarriers [4–6], nanoemulsions made of biodegradable materials are easily manufactured and result in high biocompatibility. Nanoemulsions are finely dispersed systems where nanoscale droplets of one liquid are stabilized within another, typically comprising an oil-in-water or water-in-oil structure. Due to their size (typically under 200 nm), nanoemulsions enhance drug solubility, stability and bioavailability, which is crucial for effective drug delivery to tumours. Compared to other nanocarrier systems, such as liposomes and polymeric nanoparticles, nanoemulsions offer unique advantages, including greater stability, ease of manufacturing, and a high loading capacity for lipophilic drugs [7]. For metastatic lung cancer, nanoemulsions hold special promise [8]. Their small size allows them to penetrate deeply into tumour tissues and access metastatic cells that are often challenging to reach with conventional therapies. Additionally, their ability to encapsulate therapeutic molecules, minimizing their degradation and prolonging circulation time, enhances drug concentration at tumour sites and reduces systemic side effects [9,10]. By choosing a nanoemulsion-based delivery system, like the sphingomyelin (SM) and vitamin E (VitE)-based formulations our group has developed (SNs), there is potential for improved therapeutic efficacy and patient outcomes in metastatic lung cancer. These nanoemulsions offer a versatile platform for delivering a variety of therapeutic agents, including hydrophobic drugs, biomolecules, oligonucleotides for gene therapy, and radiometals for diagnostic applications [9,11–16], which could provide targeted and sustained treatment options.

Assessing the toxicological profile of nanoparticles is crucial, even though nanostructures based on biodegradable organic materials tend to have low or no toxicity and do not accumulate in the body [17,18]. These specific SNs have been shown to exhibit low toxicity in previous assays reported in our research group. In vitro studies with SW480 and MiaPaCa-2 cells demonstrated cytotoxic concentration (IC_50_) values 10 to 20 times higher than the average values found in many other lipidic nanosystems [15]. Additional tests with stearylamine and DOTAP incorporation showed no toxic effects and confirmed the biocompatibility of the emulsions in SW480 colon cancer cells [9]. The cytotoxicity profile remained consistent when tested with SW620 and MCF7 cells, as well as with polyethylene glycol decoration [12,16]. *In vivo* studies with zebrafish and mice also showed very low toxicity and validated their potential for gene therapy development in cancer treatment [13,19].

Biocompatibility of SNs has already been established, as well as the efficacy of SNs for targeting tumors, in particular, breast [14,16,20] and lung cancer [21,22]. Lung cancer is the most diagnosed type of cancer and the leading cause of cancer-related death worldwide, with approximately 15% of patients surviving 5 years after diagnosis. In Non-Small Cell Lung Cancer (NSCLC), highly metastatic locally and in distal organs, the 5-year mean survival is lower than 5%. We have recently shown that SNs could be specifically delivered to disseminated lung cancer cells, opening new opportunities to treat metastasis (*under review publication*).

The purpose of this work was to increase the knowledge regarding their interaction of SNs with cancer cells, particularly with lung cancer cells, and the potential mechanisms underlying that could induce toxicity. Our focus encompasses exploring the potential for damage to lysosomes or mitochondria, elucidating the intracellular trajectory of these nanocarriers, and delineating their potential applications in drug delivery.

Toxicity is often associated with the production of reactive oxygen species (ROS), and interactions with cellular organelles such as the nucleus or endoplasmic reticulum may also be associated with increased toxicity [17]. Biophysical properties including size, surface area/charge and aggregation state play a key role in cell membrane damage, being the positively charged particles the most cytotoxic [23]. Conjugation of polysaccharides was also shown to enhance the accumulation of nanoparticles in the brain, liver and spleen, which correlated to their toxicity in these organs [18]. In contrast, lipid-based nanoparticles show less toxicity, although other studies with lipid nanocapsules demonstrated varying degrees of toxicity on different breast cancer cell lines, affecting signalling, and cell fate, and inducing ROS and lipid peroxidation [24].

Examining cell interaction and cell death is crucial for a thorough investigation of a specific nanocarrier, especially when considering its therapeutic use, as is the case for SNs that have already shown potential for the development of anticancer therapeutics. Thus, identifying different signaling pathways activated by nanocarriers is critical [25].

Additionally, understanding how nanocarriers interact with cellular components, navigate intracellular pathways, and reach their mitochondrial targets is essential for optimizing their design and effectiveness. This knowledge ensures that nanocarriers can efficiently deliver drugs to mitochondria, maximizing therapeutic benefits while minimizing off-target effects and toxicity [26,27]. However, challenges arise in achieving intracellular drug accumulation, particularly in mitochondria, due to protective membranes and inner membrane negative potential.

Herein, we discuss SNs potential for adenocarcinoma, assessing cytotoxicity and intracellular fate. We examined a biocompatible, biodegradable nanometric emulsion with an oily core of vitamin E stabilized by sphingomyelin on the A549 cell line, a model for lung adenocarcinoma in which the SNs have already been efficiently tested [21,22]. We compare the SNs-induced cytotoxicity between the A549 cell line and human embryonic kidney cells (HEK293) to assess if there is a cancer-specific cytotoxic effect. This investigation improves our understanding of nanocarrier toxicity profiles, enhancing their use and efficacy as targeted delivery strategies. It also opens the door to new treatment opportunities. After exploring the intracellular distribution and proving that SNs accumulate in mitochondria, we additionally investigated the potential of drug-loaded SNs to improve the delivery of resveratrol (RSV), a naturally occurring polyphenolic compound significantly present in grapes and red wine, which has been shown to have beneficial effects in human health, including cardioprotective, antioxidant and anti-inflammatory activities and shows great promise in developing novel anticancer therapies. Indeed, several preclinical studies have revealed that RSV is a potential anticancer agent due to its chemopreventive effects on three major stages of carcinogenesis, including initiation, promotion and progression [28]. RSV can induce cell death through the mitochondrial apoptotic pathway, in which mitochondria play a central role in the release of pro-apoptotic factors [29] but present challenges for clinical applications in terms of poor water solubility, chemical instability, and low bioavailability due to intestinal metabolism [30]. Exploiting the capacity of SNs and their oily core to dissolve large quantities of hydrophobic and low soluble drugs along with their mutual compatibility and ability to protect the drugs from hydrolysis and enzymatic degradation make them suitable drug delivery vectors. In this work, the potential of these RSV-loaded SNs as mitochondria-targeting agents in A549 cells was also proved.

## 2. Materials and Methods

### 2.1 Materials

Vitamin E (VitE) was obtained from Calbiochem (Merck-Milipore, Darmstadt, Germany). Sphingomyelin (SM) was purchased from Lipoid GmbH (Ludwigshafen, Germany). SM marked with Cy5 or TopFluor were obtained from Avanti Polar Lipids (Alabama, USA).

Roswell Park Memorial Institute (RPMI) medium and Dimethyl Sulfoxide (DMSO), Phosphate Buffer Saline (PBS), Trypsin and Mowiol 4-88 were acquired from Sigma-Aldrich (Darmstadt, Germany). Fetal Bovine Serum (FBS) and Dulbecco’s Modified Eagle’s Medium (DMEM) without L-Glutamine or Phenol Red were obtained from Lomza (Basilea, Switzerland). Streptomycin and penicillin were acquired from Gibco (Massachusetts, USA). Paraformaldehyde (PFA) 16% was obtained from Thermo Fisher (Massachusetts, USA). Trypan blue was purchased from Gibco (Massachusetts, USA).

Fluorescent probes such as Hoechst 33342, MitoTracker Green FM, LysoTracker Green DND, MitoProbe DiIC_1_ (5) Assay Kit for Flow Cytometry, Annexin V and propidium iodide (PI) for Flow Cytometry and alamarBlue Cell Viability Reagent, were obtained from ThermoFisher Scientific (Massachusetts, USA). Autophagy Assay Kit MAK138 was purchased from Sigma-Aldrich, Merck (Darmstadt, Germany) and 7-Amino-Actinomycin D (7-AAD) was obtained from BD Bioscences (New Jersey, USA). Resveratrol (RSV) was obtained from Cayman Chemicals (Michigan, USA).

### 2.2. Preparation and characterization of SNs

SNs made only of Vitamin E and sphingomyelin, two natural components of cell membranes, have been formulated by a simple and well-reproducible ethanol injection method as previously described (Figure 1) [15,20]. One hundred microliters of the organic phase (composed of VitE, SM and ethanol) were injected into one millilitre of ultrapure water at room temperature, in a molar ratio of VitE:SM of 1:0.1, according to previous optimization and characterization studies [15].

**Figure 1.**
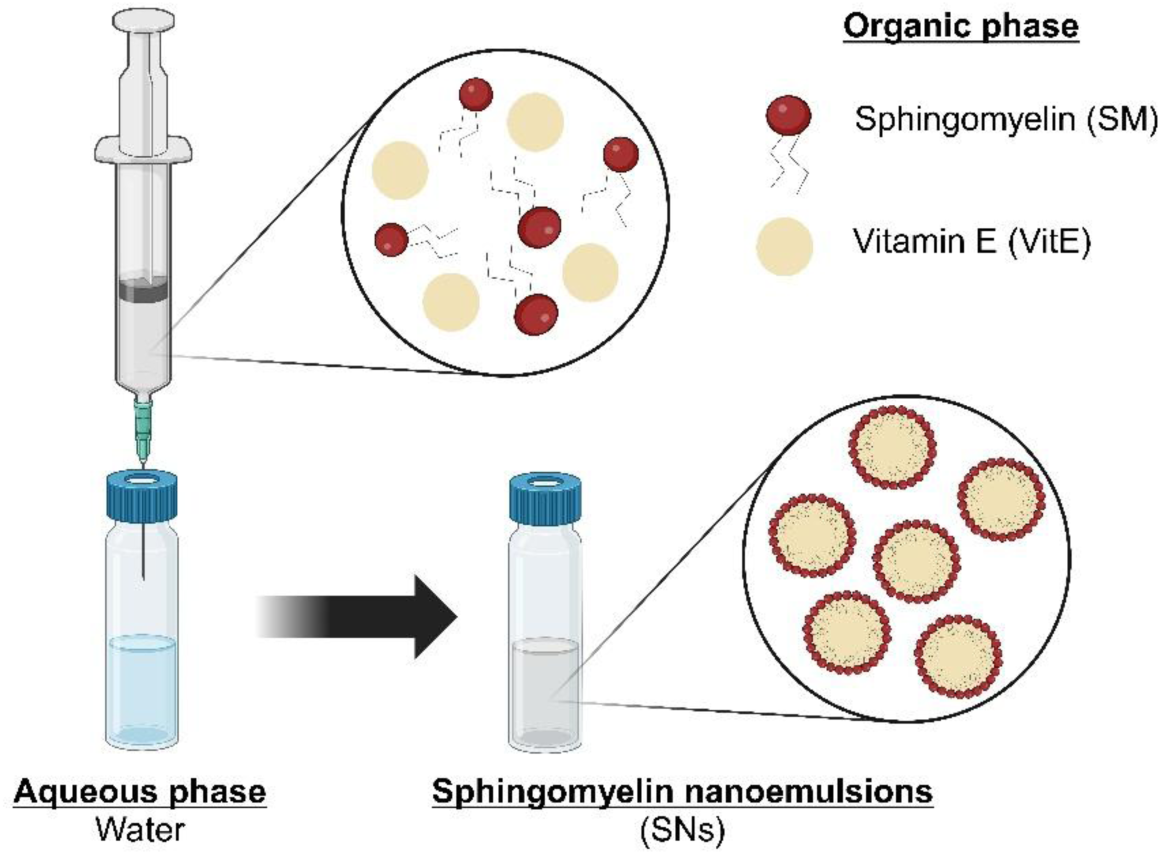
Schematic of the formulation method using ethanol injection. SNs are composed of vitamin E and sphingomyelin.

The resulting nanosystems were characterized in terms of particle size, polydispersity index (PdI), and Z-Potential by using a Zetasizer Nano ZS (Malvern Instruments Ltd., UK) to measure Dynamic Light Scattering and Laser Doppler Anemometry. For size and polydispersity index measurements, samples were diluted 1:10 (v/v) with ultrapure water and were performed at 25°C with a detection angle of 173°. For the determination of Z-potential, the formulations were further diluted 1:100 (v/v) in ultrapure water. Colloidal stability was assessed in short-term and long-term stability and relevant biological medium. Short-term stability was performed for early determination of immediate instability behaviour. Colloidal stability was also determined in a relevant biological medium, i.e. supplemented cell culture medium. Briefly, SNs were diluted 1:10 (v/v) with the correspondent medium for incubation (RPMI medium supplemented with 10% FBS) reaching a final concentration of 1 mg/mL. Nanosystems then were incubated at 37°C under constant horizontal shaking for up to 24h. For measuring purposes, the previous mixture was further diluted 1:10 (v/v) in water and analyzed by DLS.

### 2.3. Preparation of fluorescent-labelled SNs

Fluorescent-labelled nanosystems were prepared as described in the previous section ***¡Error! No se encuentra el origen de la referencia.***. SM-TopFluor was solubilized in ethanol, incorporated in the organic phase, and consequently injected into the aqueous phase for a final concentration of 4 µg/mL. In the case of C16 SM-Cy5 labelled SNs, the fluorophore is dissolved in chloroform, so it must be evaporated before its incorporation into the ethanolic phase. Then, it is injected into 1 mL of ultrapure water to get a final concentration of 2 µg/mL. SM-Cy5 and SM-TopFluor will be incorporated into the lipidic structure of the nanosystem.

### 2.4. Preparation of RSV-loaded SNs (RSV-SNs)

RSV-loaded SNs (RSV-SNs) were prepared analogously as previously described in sections ***¡Error! No se encuentra el origen de la referencia.*** and *Preparation of fluorescent-labelled SNs.* RSV was solubilized in ethanol at a concentration of 20 mg/mL in the stock solution, and incorporated in the organic phase, to consequently be injected into the aqueous phase for a final concentration of 100 µg/mL. RSV will be incorporated into the lipidic structure of the nanosystem. RSV quantification was performed using High-Performance Liquid Chromatography (HPLC) using a Reverse Phase HPLC Column Poroshell 120-C18, 4.6 Å x 100 mm. Methanol + 2% 2-propanol and MiliQ H_2_O + 2% 2-propanol were used as mobile phases. The method is based on a 10-minute run at 0.5 mL/min. A calibration curve was prepared for RSV ranging from 0 to 100 µg/mL in methanol and absorbance signal was recorded at 303 nm. RSV quantification from the SNs was performed after ultrafiltration using Amicon 10 KDa filters (Amicon Ultra-0.5 Centrifugal Filter Unit, Merck Millipore, Darmstadt, Germany). To calculate encapsulation efficiency (EE%), the following equation was used:

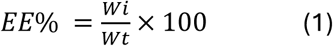

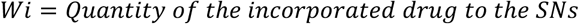

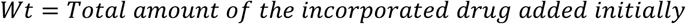

### 2.5. Cell culture

The human lung adenocarcinoma cell line (A549) was purchased from American Type Cell Culture (ATCC) and maintained according to recommendations. Briefly, A549 cells were cultured in RPMI medium supplemented with 10% heat-inactivated FBS and penicillin (50 U/mL) and streptomycin (0.05 mg/mL). The cell line was maintained in a humidified incubator at 37°C and 5% CO_2_.

The human embryonic kidney cell line (HEK293) was purchased from American Type Cell Culture (ATCC) and maintained according to recommendations. HEK293 cells was cultured in DMEM supplemented with 10% heat-inactivated FBS and penicillin (50 U/mL) and streptomycin (0.05 mg/mL). The cell line was maintained in a humidified incubator at 37°C and 5% CO_2_.

### 2.6. Cell viability assay

Cells were seeded onto 96-well plates at a density of 10,000 cells per well and incubated overnight under standard culture conditions. Then, the cell medium was replaced by SNs and cell medium at the desired concentration. After incubation with SNs, the cell medium was removed, and cells were washed with PBS. Then, cell viability was measured with alamarBlue Cell Viability Reagent (ThermoFisher Scientific, USA) according to the manufacturer’s instructions. Briefly, the reagent was diluted in a non-supplemented RPMI medium at a final 1:10 (v/v) added to cells and incubated for 3h at 37°C protected from light. Fluorescence was measured at BMG Labtech Plate Reader at 530/590 nm (excitation/emission). Non-treated cells were considered as the negative control (100% of viability) and cells treated with Triton X-100 were considered as the positive control (0% of viability). For the remaining treated cells, viability was calculated accordingly.

For IC_50_ calculation, cells were incubated for 24h with the following range of lipid concentrations: 2.750, 1.375, 0.660, 0.330 and 0.165 mg/mL. IC_50_ was calculated upon a semi-log dose-response curve. Analogously, HEK293 cells were incubated for 72h with SNs at the same range of lipid concentrations (2.750-0.165 mg/mL) and IC_50_ was calculated upon a semi-log dose-response curve.

For IC_50_ of SNs-RSV, cells were incubated for 24h with the following range of drug concentrations: 219.06, 131.43, 52.57, 26.29 and 13.14 µM. IC_50_ was calculated upon a semi-log dose-response curve.

For the determination of a non-cytotoxic range, A549 cells were incubated 24, 48 or 72h with the following range of lipid concentrations: 0.220, 0.110, 0.055, 0.028, 0.014 mg/mL.

### 2.7. Cell death study: annexin V and Propidium Iodide (PI)

A549 cells were seeded onto 6-well plates at a density of 300,000 cells per well and incubated overnight under standard culture conditions. Cells were incubated for 24h with SNs at concentrations equivalent to their IC_50_ values. Cells were harvested and washed with PBS. Then, re-centrifuge the washed cells. Alexa Fluor 488 annexin V and PI at a concentration of 100 μg/mL were added to each sample. Cells were incubated for 15 min at room temperature covered from light. Finally, samples were homogenized in 400 µL 1X annexin-binding buffer and fluorescence was analyzed with BD FACSAria IIu sorter. Alexa Fluor 488 annexin V fluorescence was monitored using an excitation filter of 488 nm and an emission wavelength of 660 nm. PI fluorescence was monitored using an excitation filter of 488 nm and an emission wavelength of 530 nm.

### 2.8 Measurement of caspases 3 and 7activation

A549 cells were seeded onto 6-well plates at a density of 300,000 cells per well and incubated overnight under standard culture conditions. Cells were incubated for 24h with SNs at concentrations equivalent to their IC_50_ values. Cells were harvested with trypsin. CellEvent Caspase-3/7 Green Detection Reagent at a concentration of 500 μM was added to each sample. Cells were incubated for 25 min at 37°C protected from light and SYTOX AADvanced at a concentration of 1 mM was added to each sample. Cells were incubated for 5 min at 37°C and protected from light. Finally, samples were homogenized, and fluorescence was analyzed with BD FACSAria IIu sorter. CellEvent Caspase 3/7 Green Detection Reagent fluorescence was monitored using an excitation filter of 511 and an emission wavelength of 533 nm. SYTOX AADvanced fluorescence was monitored using an excitation filter of 546 nm and an emission wavelength of 647 nm.

### 2.9. Determination of SNs uptake by confocal microscopy

A549 cells were seeded onto an 8-well µ-chamber at a density of 70,000 cells per well and incubated overnight under standard culture conditions. The cell medium was replaced by 0.0176 mg/mL of total lipid concentration of SNs-labelled with TopFluor diluted in RPMI supplemented with 10% FBS and penicillin/streptomycin. Cells were incubated with the SNs for 30 min, 1, 2 and 4h and then fixed with PFA 4% overnight. Samples were analyzed with a Multiphoton Microscope Leica TCS SP5 MP. TopFluor fluorescence was monitored using an excitation filter of 495 nm and an emission wavelength of 503 nm.

### 2.10. Determination of lysosomal co-localization

A549 cells were seeded at FluoroDish at 300,000 cells per well and maintained overnight under standard culture conditions. Cells were incubated for 2h protected from light with Cy5-labelled SNs at concentrations equivalent to their IC_50_ values. Cells were washed once with PBS and nuclei were stained with Hoechst 33342 (stock dilution 10 mg/mL was diluted 1:1000 in PBS). After 15 min incubation protected from light at room temperature, 1 μL LysoTracker Green DND 5·10^4^ nM was added to each well and fluorescence was analyzed immediately with Multiphoton Microscope Leica TCS SP5 MP. Hoechst 33342 fluorescence was monitored using an excitation filter of 460 nm and an emission wavelength of 350 nm. LysoTracker Green FM fluorescence was monitored using an excitation filter of 504 nm and an emission wavelength of 511 nm. Cy5 fluorescence was monitored using an excitation filter of 647 nm and an emission wavelength of 665 nm.

### 2.11. Analysis of acidic lysosome

A549 cells were seeded at 6-well plates at 300,000 cells per well and incubated overnight under standard culture conditions. Then, cells were treated with SNs at concentrations equivalent to their IC_50_ values for 24h. Cells were harvested and washed with PBS. LysoTracker Green DND 5·10^4^ nM was added to each sample. samples were homogenized in 500 µL iced-cold PBS and fluorescence was analyzed with BD FACSAria IIu sorter. LysoTracker Green DND was monitored using an excitation filter of 504 nm and an emission wavelength of 511 nm.

### 2.12. Analysis of autophagosome formation

Cells were seeded onto 96-well plates at a density of 10,000 cells per well and incubated overnight under standard culture conditions. Then, cells were treated with SNs at concentrations equivalent to their IC_50_ values for 24h. Autophagosome Detection Reagent 500x was added to each sample. Cells were incubated for 30 min at 37°C protected from light and washed twice with PBS. Finally, the fluorescence is read on the plate reader VICTOR Nivo using an excitation filter of 360 nm and an emission wavelength of 520 nm.

### 2.13. Measurement of mitochondrial co-localization

A549 cells were seeded at FluoroDish at 300,000 cells per well and maintained overnight under standard culture conditions. Cells were incubated with Cy5-labelled SNs at concentrations equivalent to their IC_50_ values, previously determined. Treated cells were incubated for 2h protected from light. Cells were washed with PBS and nuclei were stained with Hoechst 33342 (stock dilution 10 mg/mL was diluted 1:1000 in PBS). After 15 min incubation in the dark at room temperature, cells were washed with PBS and then MitoTracker Green FM solved in non-supplemented RPMI medium at 150 nM was added to each well. Cells were incubated for 20 min and fluorescence was analyzed with Multiphoton Microscope Leica TCS SP5 MP. Hoechst 33342 fluorescence was monitored using an excitation filter of 460 nm and an emission wavelength of 350 nm. MitoTracker Green FMfluorescence was monitored using an excitation filter of 490 nm and an emission wavelength of 516 nm. Cy5 fluorescence was monitored using an excitation filter of 647 nm and an emission wavelength of 665 nm.

### 2.14. Analysis of mitochondrial membrane potential (ψ_m_) integrity

A549 cells were seeded at 6-well plates at 300,000 cells per well and maintained overnight under standard culture conditions. Then, cells were treated with SNs at concentrations equivalent to their IC_50_ values for 24h. Cells were harvested and washed with PBS. DiIC_1_ diluted in DMSO at a concentration of 10 µM was added to each sample. Cells were incubated for 30 min at 37°C covered from light and washed twice with PBS. Finally, samples were homogenized in 500 µL iced-cold PBS and fluorescence was analyzed with BD FACSAria IIu sorter. DiIC_1_ fluorescence was monitored using an excitation filter of 638 nm and an emission wavelength of 658 nm.

### 2.15. Cell uptake by energy-dependent pathways determination

A549 cells were seeded at two 6-well plates at a density of 300,000 cells per well and maintained overnight under standard culture conditions. Then, one plate was incubated for 30 min at 4°C, whereas the other was maintained at 37°C. After that, the cell medium was replaced by SNs at the desired concentration. Cell medium without SNs was added for negative control. After 2h incubation at 4°C or 37°C, cells were harvested and washed twice with PBS. Finally, samples were analyzed with BD FACSAria IIu sorter. TopFluor fluorescence was monitored using an excitation filter of 495 nm and an emission wavelength of 503 nm.

### 2.16. Statistical analysis

Results were shown as mean ± SD. Firstly, the Shapiro-Wilk test, a normality test for small samples (n<30), was performed to observe whether the data followed a normal distribution or not. The significance of differences between two groups was determined via the t-student test, and the significance of differences between more groups was determined via one-way analysis of variance (ANOVA). All statistics were conducted using GraphPad Prism 8.0, and differences were considered significant if p<0.05 (*), p<0.01 (**), p<0.001 (***), p<0.0001 (****).

## 3. Results

This section may be divided by subheadings. It should provide a concise and precise description of the experimental results, their interpretation, as well as the experimental conclusions that can be drawn.

### 3.1. Preparation and Characterization of SNs

SNs were prepared and characterized as previously reported [22], and results show a mean size of 113 nm, a polydispersity index (PdI) of 0.10 and a zeta potential of −11.8 mV, in agreement with previously cited works [16,31]. The PdI, a critical parameter reflecting nanoparticle uniformity, was 0.1 throughout the study, indicating a highly monodisperse nature (values ≤0.1 are considered optimal).

Next, the assessment of SNs stability was examined in RPMI medium supplemented with FBS. The results illustrated in Figure 2 confirm the stability of SNs in RPMI medium supplemented with FBS. No significant differences in size or PdI were observed, underscoring the robustness of the nanocarrier under the given conditions.

**Figure 2.**
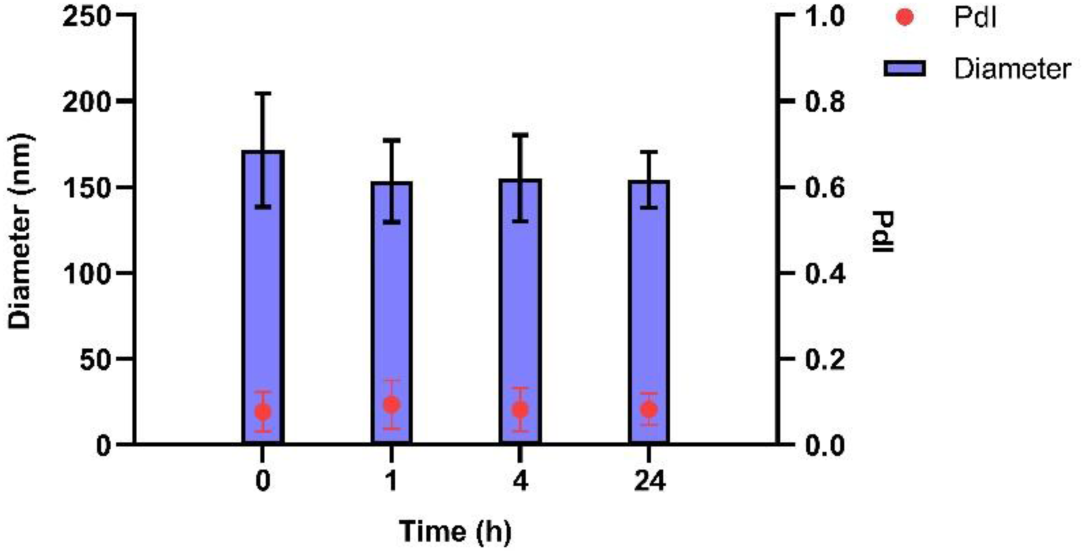
Measurement of the stability of SNs in RPMI medium supplemented with 10% FBS and penicillin/streptomycin. Changes in diameter (nm) and PdI value over time are shown. Results are expressed against negative control (culture medium) as mean ± standard deviation of at least three independent assays.

### 3.2. Cytotoxicity of SNs in the A549 cell line

The cytotoxic effect of SNs was assessed using the alamarBlue assay. After 24 and 72 hours of incubation with SNs, the IC_50_ values for the A549 cell line were calculated as 0.89 mg/mL and 0.63 mg/mL, respectively (Figure 3). These results demonstrated a concentration- and time-dependent cytotoxicity. Comparatively, the IC_50_ values for the HEK293 cell line after 72 hours of incubation (Figure 4) were 1.133 mg/mL. This value indicates that SNs are almost twice as cytotoxic to the tumor cell line compared to the normal cell line. To corroborate this, the selectivity index (SI) was calculated to quantify the selectivity of the SNs for cancer cells over normal cells using the equation:

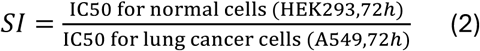

**Figure 3.**
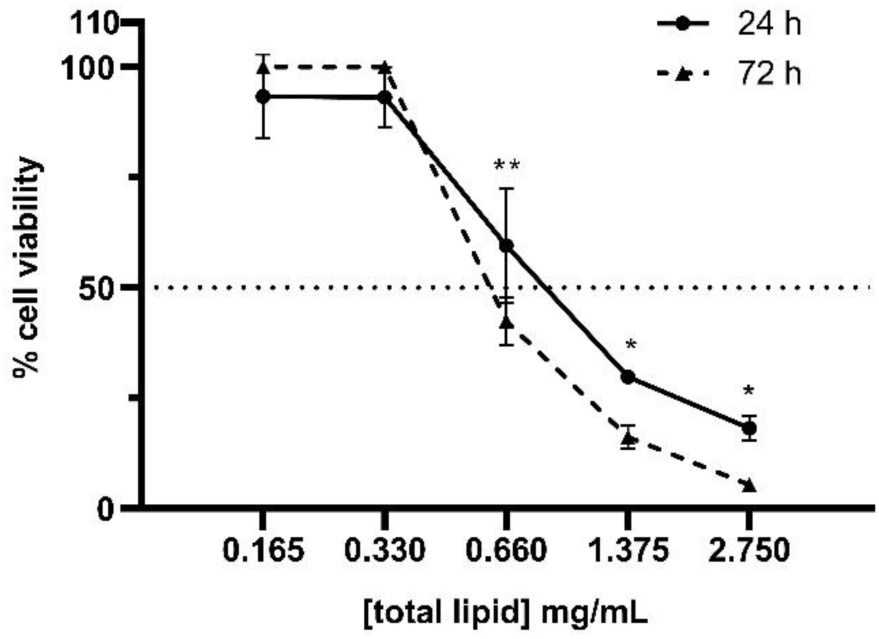
Percentage of A549 cell viability upon 24 or 72h incubation with SNs at 2.750-0.165 mg/mL of total lipid. IC_50_ value at 24h incubation is 0.89 ± 0.15 mg/mL, at 72h incubation is 0.63 ± 0.03 mg/mL. Data are expressed as mean ± standard deviation of four independent assays. *: p<0.05. **: p<0.01.

**Figure 4.**
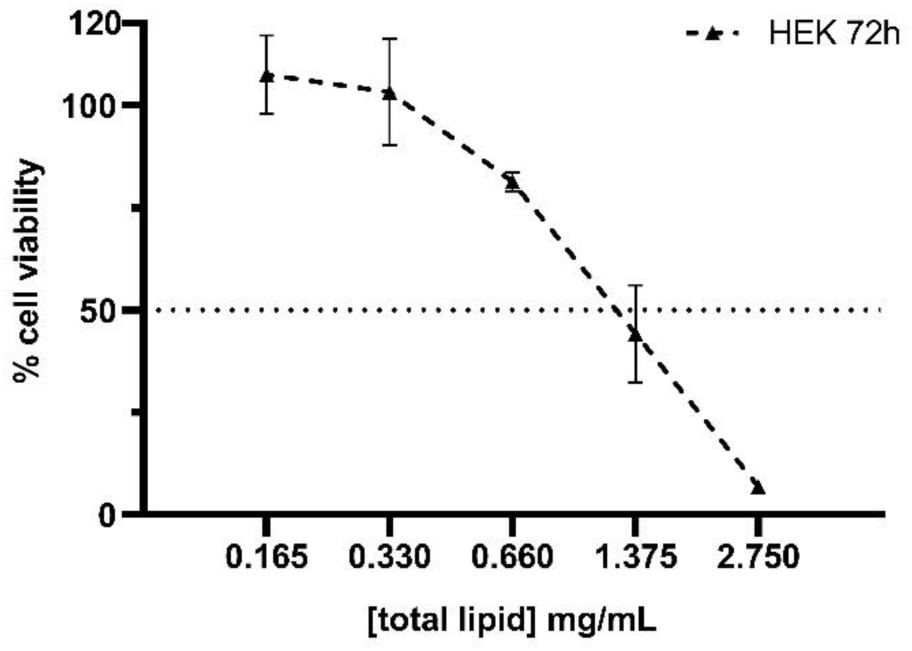
Percentage of HEK293 cell viability upon 72h incubation with SNs at 2.750-0.165 mg/mL of total lipid. IC_50_ value at 72h incubation is 1.133 ± 0.066 mg/mL. Data are expressed as mean ± standard deviation of four independent assays.

A value of 1.8 was obtained, indicating greater selectivity for cancer cells.

### 3.3. Mechanisms of cell death

Annexin V and PI assays revealed a significant increase in apoptotic A549 cells after 24 hours of incubation with the IC_50_ concentration of SNs (Figure 5A). Around 20% of cells were detected in early and late apoptosis stages. Further validation using caspase-3/7 assays confirmed the activation of apoptotic pathways. The activation of caspases 3/7 was measured in cells treated with IC_50_ dose of SNs after 24 h incubation (Figure 5B). The levels of caspase expression were compared with untreated cells. Although an 80% cell population was detected after treatment with SNs, this does not imply that 80% of cells are alive; rather, it suggests that these cells lack activated caspases 3 and 7.

**Figure 5.**
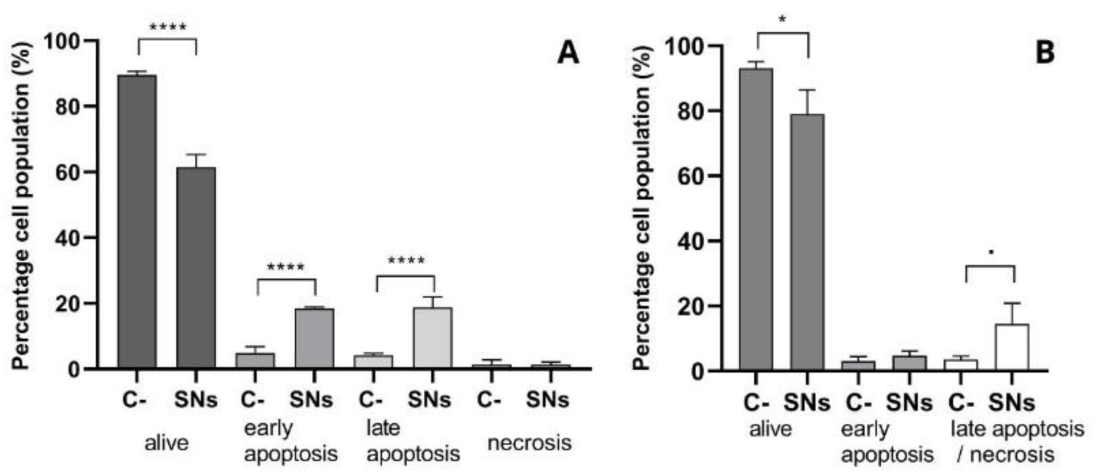
Percentage of the cell population of A549, after 24h incubation with RPMI supplemented with 10% FBS and penicillin/streptomycin (C-), SNs at a concentration equivalent to their IC_50_. (A) Using Annexin V and PI assay by flow cytometry. The early apoptosis (Annexin V^+^, 7-PI^−^), the late apoptosis (Annexin V^+^, PI^+^), the alive cells (Annexin V^−^, PI^−^), the necrosis (Annexin V^−^, PI^+^). Results are expressed against negative control (culture medium) as mean ± standard deviation of at least three independent assays. (B) Determined with CellEvent Caspase-3/7 Green Detection Reagent and SYTOX AADvanced assay by flow cytometry. Percentage of viable A549 cells, in early apoptosis or late apoptosis/necrosis. **·**: p<0.1 *: p<0.05. ****: p<0.0001.

### 3.4. Intracellular trafficking and cell uptake by energy-dependent pathways determination

Cellular uptake kinetics were analyzed by incubating A549 cells with TopFluor-labelled SNs for different time intervals (30 min, 1, 2 and 4h) (Figure 6). Intracellular fluorescence was observed after just 30 min of incubation (Figure 6A), indicating rapid cellular uptake. Furthermore, SNs demonstrated active, energy-dependent entry into A549 cells, as shown by assays measuring cell uptake of SNs after 30 minutes of incubation at 4°C or 37°C (Table 1).

**Figure 6.**
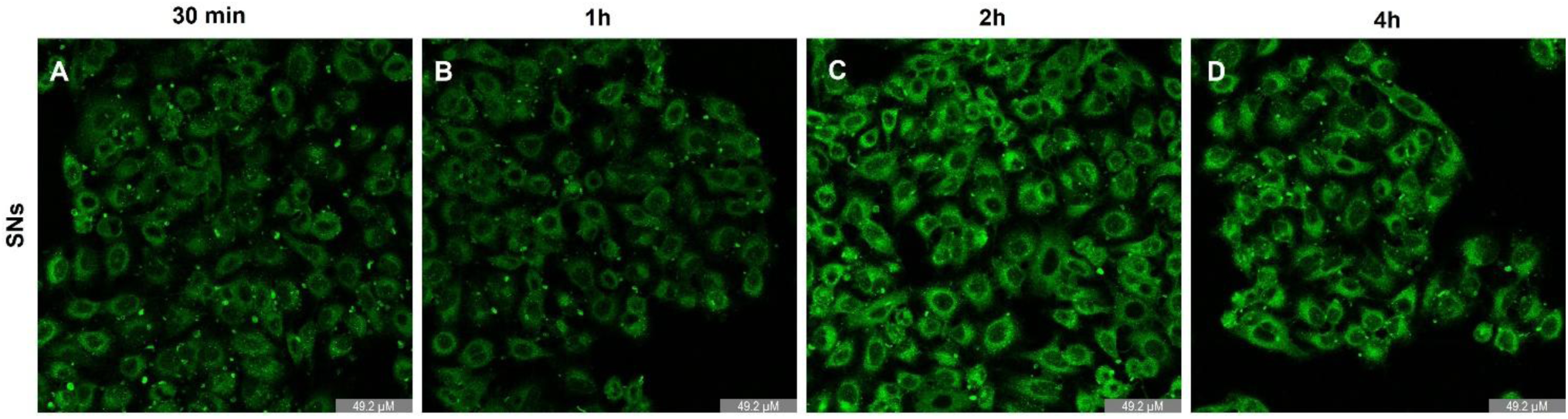
A549 incubated with non-toxic concentrations (0.0176 mg/mL of total lipid) of the SNs (green) for 30 min (A), 1h (B), 2h (C) or 4h (D). Results of three independent assays.

**Table 1.** Cell uptake of SNs on A549 cell line upon 30 min incubation at 4°C or 37°C to determine whether SNs entrance was energy-dependent. Results are expressed as percentage of fluorescent cells, indicative of SNs entrance. Negative control: untreated cells. Data are expressed as mean ± standard deviation of three independent assays.

| Condition | Control | 4°C | 37°C |
| --- | --- | --- | --- |
| Fluorescent cells (%) | 0.63 ± 0.42 | 0.53 ± 0.35 | 45.37 ± 17.10 |

To evaluate the presence of SNs in A549 lysosomes, fluorescence microscopy assays were conducted after 2h and 24h of incubation, as shown in Figure 7 (A to F). Results indicate that SNs accumulate in lysosomes (Figure 7B, C, E, F). Image analysis using Pearson’s co-localization test, which assesses the correlation of intensity distribution between channels, yielded values of 0.39 after 2h of incubation and 0.54 after 24h (+1 indicate perfect co-localization, while values of −1 indicate reverse co-localization)[32].

**Figure 7.**
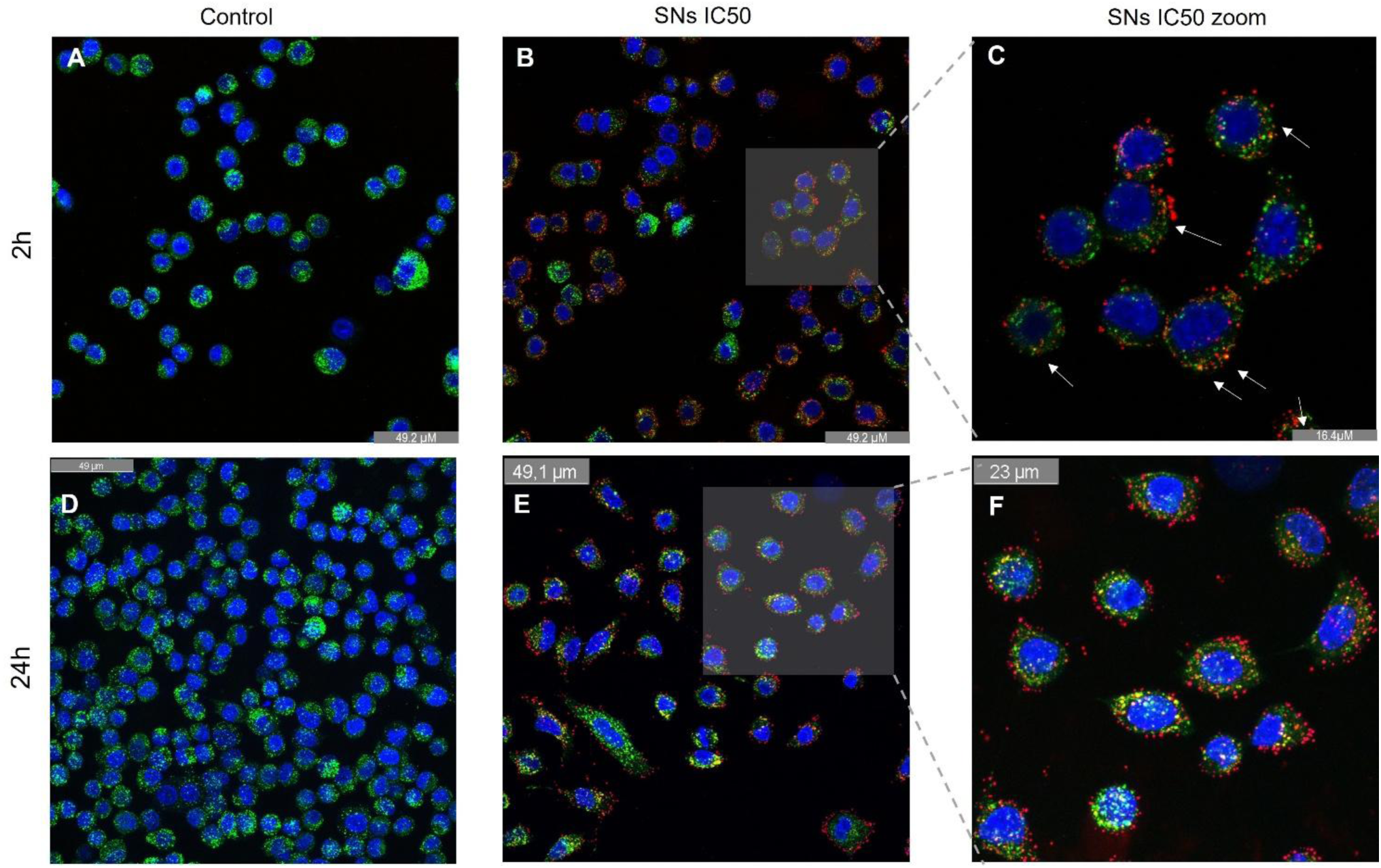
Effect of SNs on the lysosome of A549 cells after 2 and 24h incubation. Confocal microscopy images (63x) showing the staining of nuclei in blue and lysosome in green of A549 cells incubated with medium upon 2h (A) and 24h incubation (D), and with cytotoxic concentrations (IC_50_) of SNs (red) after 2h (B) and 24h of incubation (E). Zoom x3 after 2h of incubation (C) and zoom x1.6 after 24h of incubation (F). Co-localization of lysosome and nanoparticles is marked with white arrows. Results of three independent assays.

Lysosomal integrity in response to incubation with the SNs was also analyzed. The loss of lysosomal acidification observed after treatment with SNs, as measured by the decrease in functional, acidic lysosomes using Lysotracker Green and 7-AAD after 24 hours of incubation, suggests compromised lysosomal function. Autophagosome detection revealed a decrease in autophagic flux following lysosomal alterations (Figure 8).

**Figure 8.**
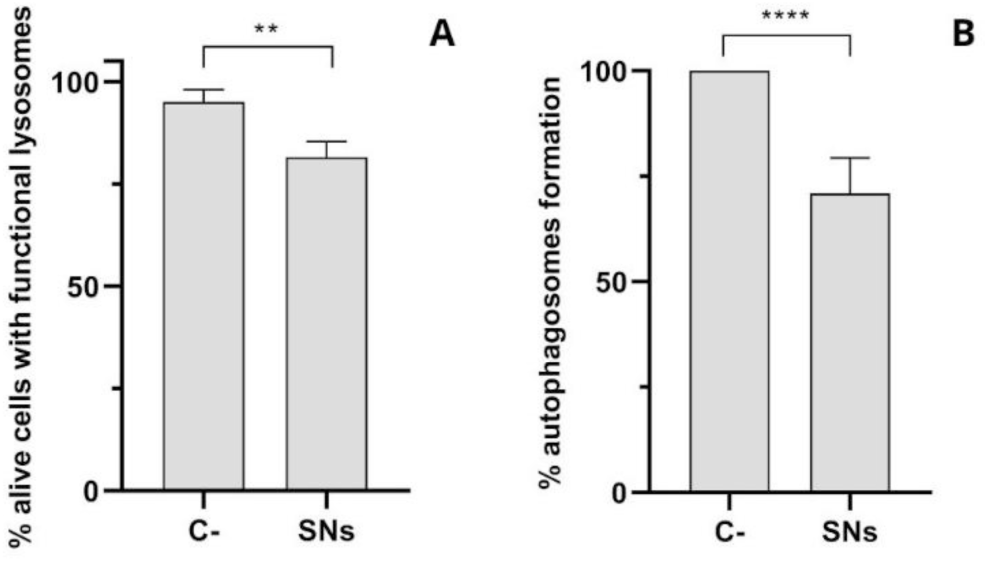
(A) Percentage of alive cells with functional lysosomes in A549 cells after 24h incubation with SNs at a concentration equivalent to their IC_50_, determined with Lysotracker Green and 7-AAD assay by flow cytometry. Results are expressed against negative control (culture medium) as mean ± standard. **: p<0.01. Results of five independent assays. (B) Percentage of autophagosome formation on A549 cells after 24h incubation with SNs, at a concentration equivalent to their IC_50_, determined with Autophagosome Detection Reagent assay by plate reader. Results are expressed against negative control (culture medium) as mean ± standard. ****: p<0.0001. Results of five independent assays.

### 3.5 Mitochondrial impact

We evaluated the capacity of these SNs to internalize into mitochondria after 2 and 24h of incubation, as shown in Figure 9. Image analysis using Pearson’s co-localization test revealed a value of 0.53 in mitochondria after 2h incubation and 0.75 after 24h incubation, indicating approximately 75% co-localization in this organelle. We further analyzed mitochondrial membrane potential (ΔΨ_m_), strongly related to mitochondrial integrity and function, to assess whether SNs internalization caused mitochondrial damage. As shown in Figure 10, significant changes in ΔΨ_m_ were observed after 24h of incubation with IC_50_ of SNs, suggesting mitochondrial damage.

**Figure 9.**
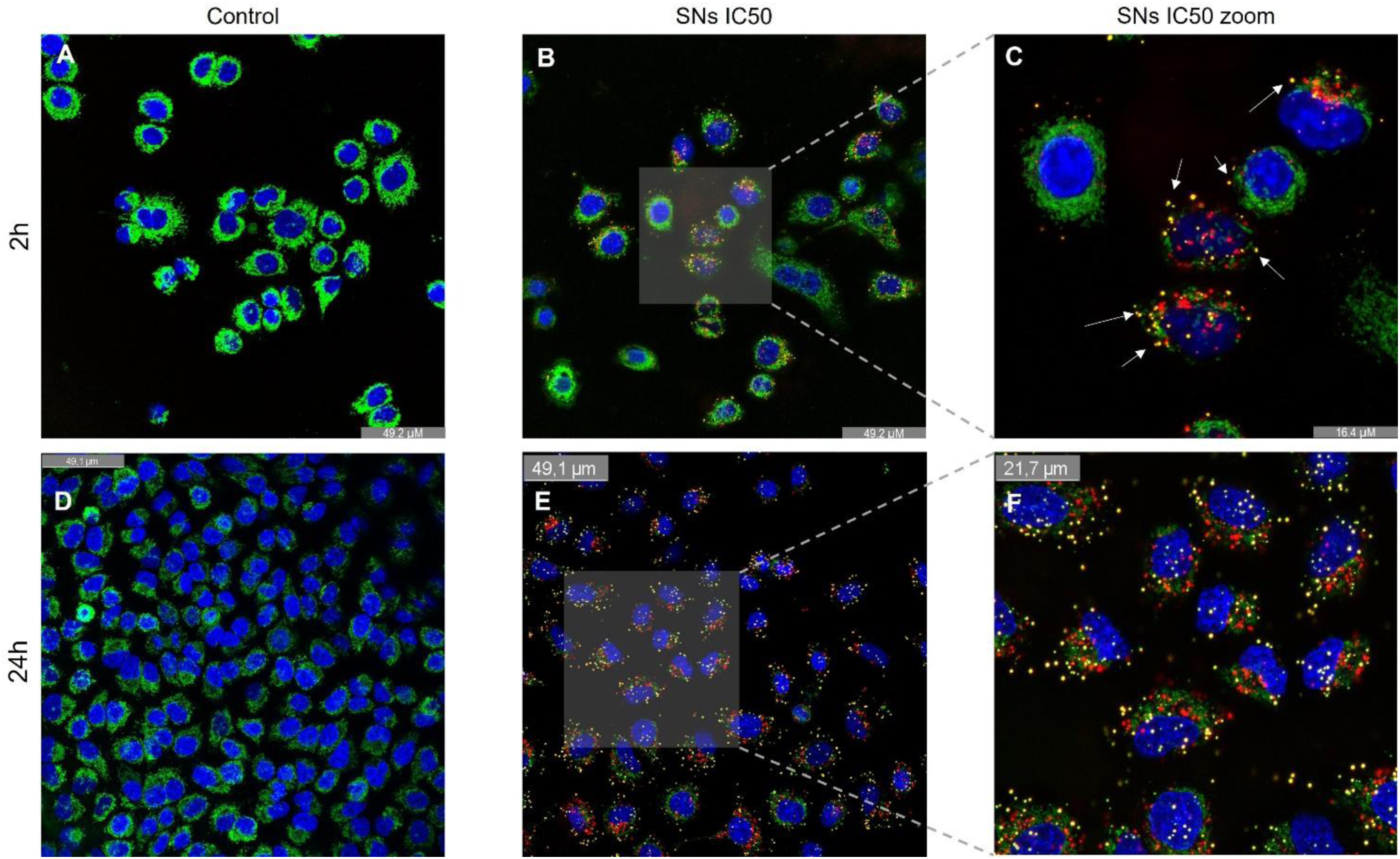
Internalization of SNs on the mitochondria of A549 cells. Confocal microscopy images (63x) showing the staining of nuclei in blue and mitochondria in green with cytotoxic concentrations of SNs (red) upon 2h (B) and 24h (E) of incubation. Co-localization of mitochondria and nanoparticles is marked with white arrows. Zoom x3 after 2h of incubation (C) and zoom x1.7 after 24h of incubation (F). Results of five independent assays.

**Figure 10.**
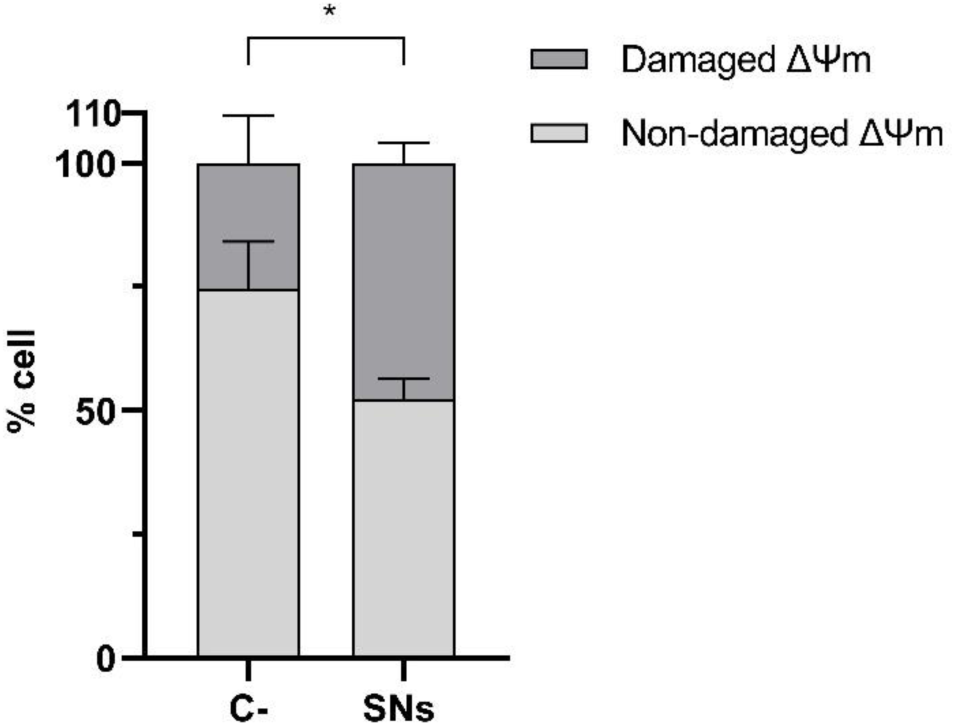
Effect of SNs on the mitochondrial membrane potential (ΔΨ_m_) of A549 cells. The percentage of total cells with damaged and non-damaged ΔΨ_m_ upon 24h incubation with cytotoxic concentrations (IC_50_) of SNs are shown. Results are as mean ± standard deviation of at least three independent assays. *: p<0.05.

### 3.5. RSV encapsulation in SNs

Encapsulation efficiency (EE%) for RSV in SNs was calculated as 82% according to equation (1). Encapsulated RSV showed improved cytotoxicity with a reduced IC_50_ value of 85.7 µg/mL compared to 118.5 µg/mL for the free drug (Table 2).

**Table 2.** IC_50_ values (µg/mL) of the drug on A549 cell lines calculated upon 24h incubation. Data are expressed as mean ± standard deviation of three independent assays.

| Condition | RSV<br>(free drug) | SNs-RSV<br>(nanoencapsulated) |
| --- | --- | --- |
| $IC_{50}$ | 118.54 ± 5.14 | 85.74 ± 4.01 |

## 4. Discussion

The findings confirm that SNs exhibit promising characteristics as nanocarriers, including a highly monodisperse nature and stability in biologically relevant conditions. The stability observed in RPMI medium with FBS underscores the potential for SNs in drug delivery applications.

Although the cytotoxicity values expected for lipid nanoparticles considered non-toxic vary depending on the specific formulation and the cell type, these data show relatively low toxicity compared to other similar nanosystems. For example, a study on lipid-based nanocarriers, including liposomes and siRNA-loaded lipid nanoparticles (LNP-siRNA) showed toxicity in HL60 and A549 cells at concentrations above 128 and 16 µg/mL, respectively [33]. Another study on cationic solid lipid nanoparticles deemed them non-toxic with IC_50_ values from 300 to 900 µg/mL [34]. Additionally, research on lipid nanocapsules found that these nanoparticles could be toxic at high concentrations with cell viability dropping below 20% across all cell lines after incubation with 1 mg/mL for 2.5 h [35]. Another example using PEGylated lipid nanocapsules as anti-cancer drug delivery systems reported IC_50_ values of 10-20 µg/mL, depending on the cell line [36].

These examples demonstrate that the obtained cytotoxicity values for SNs are among the lowest in comparison to similar lipid nanoformulations considered safe and non-toxic carriers for drug delivery.

The cytotoxicity results highlight the selective toxicity of SNs, with a favorable SI of 1.8, suggesting preferential targeting of cancer cells over normal cells. This specificity may result from increased uptake of SNs in cancer cells due to their altered lipid metabolism, as supported by previous studies. This suggests that SNs preferentially induce cytotoxicity in lung cancer cells compared to normal cells, highlighting the potential of SNs to provide more efficient targeted treatment while minimizing harm to healthy tissues. A favourable SI value is generally considered to be greater than 1.0 [37].

This specific anti-cancer effect of SNs on human lung cancer (A549) could be due to a higher uptake of SNs in cancer cells than in normal ones. This is consistent with other studies reported for the same cell line, where A549 cells also showed higher induced cytotoxicity as a consequence of enhanced cellular uptake of hydroxyapatite nanoparticles than normal bronchial epithelial cells (16HBE)[38]. Further research is needed to investigate the underlying mechanisms of SN-induced selective cytotoxicity to achieve better outcomes for clinical applications.

Apoptosis plays a vital role in maintaining homeostasis and is characterized by several morphological and biochemical changes in cells. Mechanistic studies of cell death confirmed apoptosis as the primary cytotoxic pathway, with early and late apoptosis stages implicated. Early apoptosis is characterized by cell shrinkage, membrane blebbing, and the exposure of “eat-me” signals, facilitating apoptotic cell clearance. This mechanism is preferable for targeted drug delivery to eliminate cancer cells while minimizing inflammation and immune response. Late apoptosis, also known as secondary necrosis, occurs when the plasma membrane becomes permeabilized, leading to the release of immunostimulatory molecules and autoantigens to the immune system [39], which could trigger an immune response against cancer cells. In this case, approximately 20% of the cell population was detected in both early and late apoptosis stages in A549 cells treated with IC_50_ concentration of SNs after 24h of exposure. However, care should be taken to ensure that Annexin V^+^ is indeed dying since some researchers have suggested that Annexin V can bind healthy cells that have the potential to continue proliferating. Conversely, some cells may not bind Annexin V when alive or dead. These potential issues can be overcome by following the assay to its completion (all cells PI^+^) or by measuring other features of cell death, such as caspase 3 activity [40]. Thus, these results were further validated with CellEvent Caspase-3/7 Green Detection Reagent and SYTOX AADvanced assay, which estimates the activation of executioner caspases 3 and 7 during the final stages of apoptosis.

Caspase-3 is a key mediator of apoptosis involved in the execution of cell death and has been shown to promote genetic instability and carcinogenesis when activated [41]. Additionally, caspase-3 activities can affect the survival, proliferation, and differentiation of both normal and malignant cells and tissues. Caspase-7, on the other hand, has distinct roles during apoptosis, including promoting ROS production and cell detachment, which may aid in the removal of apoptotic cells from the microenvironment [42]. Caspase activation underscores the potential for SNs to induce controlled apoptotic pathways, reducing inflammation and enhancing targeted effects.

Therefore, the activation of caspase-3 and caspase-7 by a nanocarrier may have different effects on cell fate and the surrounding microenvironment. While caspase-3 activation may lead to cell death and genetic instability, caspase-7 activation may contribute to ROS production and the detachment of apoptotic cells. Upregulation of these apoptotic mediators was observed after exposing A549 cells to SNs. Differences between the early apoptosis population were also detected between both assays, likely due to the limitations inherent to each assay. The Annexin V assay may encounter challenges in interpreting results after mechanical disaggregation of tissues, enzymatic or mechanical detachment of adherent cells, or cell transfection, which can influence phosphatidylserine (PS) flipping and introduce experimental bias. In contrast, the caspase 3/7 assay measures the activity of executioner proteases involved in apoptosis but does not directly measure the externalization of PS, a specific marker of early apoptosis detected by Annexin V.

Consequently, discrepancies arise between assays due to the different markers they detect and the complexities of the apoptotic process. While the direct effect of SNs on the activation of caspases 3 and 7 is not explicitly addressed in previous works, the broader context of caspase activation in cell death pathways and its potential modulation by sphingomyelin metabolism suggests a complex interplay that warrants further investigation in the specific context of SNs and their cytotoxic effects [43]. Overall, these findings point up the complexity of apoptosis and demonstrate that SNs incubation triggers apoptotic cell death in A549 cells, suggesting their potential for targeted cancer therapy while highlighting the necessity for further investigation into the specific pathways involved.

The mechanism by which nanostructures enter cells is largely determined by the physical and interfacial properties of nanoparticles, their interactions with the biological environment, and the properties of the cell membrane. The size, shape, and surface properties (particularly charge and hydrophobicity) of nanosystems can significantly influence cellular uptake pathways and determines their intracellular fate in terms of subcellular location and behaviour as drug nanocarriers [44,45]. In this study, we investigated the cellular uptake of the SNs in the A549 cell line. Intracellular trafficking results elucidated SNs’ rapid uptake and point out an enhancement in bioavailability, cellular permeability, and cytotoxic effects of potential encapsulated drugs in the SNs. Therefore, the quick internalization within 30 minutes highlights SNs’ great potential for efficient drug delivery. The increasing fluorescence signal with longer incubation times indicates that cellular uptake is a time-dependent process, in agreement with previous reports in the field [14]. These results align well with previous data obtained by our group in other tumor cell lines, such as the SW620 metastatic colorectal cancer cell line [12] and the breast adenocarcinoma MCF-7 cell line [16]. This suggests that the formulation leads to a significant internalization capacity in cancer cells, irrespective of the time of tumor. According to Carmona-Ule *et al*, the fast cell uptake observed might result from the enhanced lipid avidity of cancer cells due to their altered lipid metabolism, which increases the uptake of exogenous lipids [46]. Additionally, the lipidic nature of sphingomyelin, a component of cell membranes, may facilitate the interaction and uptake of these SNs by cells [47].

In addition, SNs’ energy-dependent entry mechanism (Table 1) influences the efficiency and kinetics of internalization, potentially affecting the pharmacokinetics and bioavailability of drugs or therapeutic agents delivered by SNs. This may have implications for the transport of therapeutic agents across biological barriers by influencing the mechanisms involved in cellular internalization.

Given the energy-dependent transport mechanisms through which SNs enter A549 cells, as discussed previously, it was anticipated that one or more endocytic pathways, mostly energy-dependent, could be involved. These pathways typically culminate in lysosomes, responsible for degrading endocytosed materials [48,49]. Consequently, the efficacy of nanoparticle-based therapies might be hampered by lysosomal enzyme degradation.

The lysosomal accumulation (39% of SNs co-localizing at 2h and 54% after 24h) indicates that SNs internalization into lysosomes is a time-dependent process. The Pearson correlation index can also be used to study lysosomal escape. As mentioned in the previous section (section 3.4), the coefficient varies from 0, indicating complete lysosomal escape, to 1 indicating no lysosomal escape [50]. One implication is that it may affect the release of SNs (and their potential cargo) from the lysosomes into the cytoplasm. This could impact cellular processes regulated by sphingomyelin and related molecules, such as signalling pathways and membrane homeostasis. Additionally, the time-dependent nature of lysosomal escape may influence the extent of lysosomal damage and membrane permeabilization, which can in turn affect cell viability and function. Kinetics of lysosomal escape are crucial to be well understood for the design and optimization of SNs-based drug delivery systems. The loss of lysosomal acidification observed could be attributed to the presence of SNs in the lysosome, as nanoparticle accumulation in lysosomes has been reported to adversely affect their function [52]. Additionally, the composition of SNs, resembling cell membranes due to lipid content, may facilitate lysosomal escape [51] (Figure 8A). Our findings suggest that SNs impact lysosomal pH and function, potentially leading to release into the cytoplasm, facilitated by the membrane-like composition of SNs. This could result in enhanced drug release, as toxicity to lysosomes causing lysosomal membrane permeability can trigger the release of encapsulated drugs or therapeutic agents from nanoparticles, allowing sustained and controlled release over time. Lysosomal integrity and function are also closely linked to autophagy. Lysosomal alterations might disrupt autophagic flux by inhibiting lysosome-autophagosome fusion, compromising autophagosome degradation [44,45]. This disruption has been associated with impaired cell function and cell death. Given our earlier findings of compromised lysosomal function (Figures 7E, 8A), we proceeded to assess autophagosome formation after 24 hours of incubation with Autophagosome Detection Reagent. The analysis revealed a significant decrease in autophagosome formation compared to the control (Figure 8B), suggesting an interruption in the autophagic flux. This interruption might be linked to SNs’ capacity to trigger apoptotic cell death (Figure 5), showing the role of dysfunctional autophagy in tumor progression and the potential for regulating this process to enhance the sensitivity and efficacy of cancer therapies [52,53].

Previous studies have suggested that sphingomyelin-containing liposomes can enter mitochondria via membrane fusion, primarily through micropinocytosis [54]. Once inside the cells, these liposomes escape from endosomes and subsequently penetrate mitochondria [55]. Given that SNs are lipid-based nanostructures enriched in sphingomyelin and considering the potential release of the nanoemulsion into the cytoplasm via endosomal escape due to lysosomal damage (Figure 7, Figure 8A), we hypothesized that SNs might internalize into mitochondria. Moreover, evidence suggests that sphingomyelin accumulation in mitochondria is associated with reperfusion damage, and sphingolipid dysregulation interferes with mitochondrial regulation, leading to dysfunction and cell death [56]. By targeting mitochondrial metabolism, it is possible to disrupt energy production and biosynthetic pathways crucial for cancer cell survival, making mitochondrial targeting important in diseases like cancer or metabolic diseases [57,58]. By looking at the results (Figure 9), SNs’ entry into mitochondria was a time-dependent process and is consistent with lysosomal uptake observations (Figure 7). Image analysis using Pearson’s co-localization test revealed values of 53% co-localization after 2h and 75% after 24h of incubation in this organelle. Interestingly, only 54% of SNs co-localized in lysosomes after 24h of incubation (Figure 7E). This could be attributed to lysosomal leakage caused by rupture of the lysosomal membrane at cytotoxic concentration, enabling SNs to internalize into mitochondria. These results suggest that SNs can accumulate in mitochondria in a time-dependent manner, indicating a dynamic interplay between mitochondria and lysosomes, and the potential accumulation of SNs in the mitochondria over time. This aligns with literature indicating that sphingolipids, including sphingomyelin, can impact mitochondrial function and contribute to processes like mitochondrial dynamics, mitophagy, and cell death, as well as the dynamic inter-organelle transfer of metabolites between mitochondria and lysosomes in disease pathogenesis [59–61].

Concerning the significant changes in ΔΨ_m_ observed after 24h of incubation with IC_50_ of SNs, this could indicate that SNs internalization into mitochondria may affect their function, consistent with similar findings reported in other studies with lipid nanoemulsions in A549 cells [62,63].

In summary, our results indicate that SNs can internalize or attach to mitochondria following endosomal escape, leading to modifications in mitochondrial membrane potential, indicative of dysfunction. These findings are significant as targeting mitochondria could offer potential therapeutic benefits, particularly in inhibiting cell proliferation in cancer cell lines [64] and emphasize the importance of endosomal escape and lysosomal interactions in optimizing SNs-based delivery systems. Understanding the impact of SNs on ΔΨ_m_ is relevant for the development of potential therapeutic applications.

The ability of SNs to penetrate mitochondria (Figure 8) suggests their potential as carriers for drugs targeting this organelle, such as the bioactive compound RSV [65,66]. Recently, several *in vitro* studies have the potential of RSV in managing NSCLC. Some researchers have even suggested that RSV could sensitize NSCLC cell models to conventional chemotherapeutic agents like paclitaxel [67] or etoposide [68]. Despite promising *in vivo* outcomes, achieving the concentrations required to induce apoptosis *in vivo* remains challenging [69]. Hence, encapsulating RSV in nanoparticles has emerged as a promising strategy to enhance its *in vivo* antitumor effect [70].

Using the equation outlined in the methodology section (Equation 1), we determined an average EE% of 82% for SNs. These results align with previously reported encapsulation efficiencies for mesoporous silica nanoparticles, ranging from 67% to over 93% depending on the specific formulation and encapsulation method used [71,72].

We calculated the IC_50_ values of SNs-RSV and RSV alone, as shown in Table 2. Encapsulation reduced the IC_50_ from 118.5 µg/mL (free drug) to 85.7 µg/mL (encapsulated drug), indicating a decrease in the required dose. This reduction in the IC_50_ value of SNs-RSV compared to RSV alone can be attributed to the improved accumulation into mitochondria and targeted delivery of RSV achieved through encapsulation in SNs. This facilitates increased cellular uptake and improved intracellular release of RSV, thereby enhancing its pharmacological activity. Additionally, the presumably sustained release of RSV from SNs prolongs its exposure to target cells, further enhancing efficacy on mitochondrial delivery. This aligns with the increasing focus on mitochondrial targeting in cancer therapy, leveraging SNs’ lipid composition and bioavailability.

Future studies should delve deeper into the molecular mechanisms of SNs’ interactions with lysosomes and mitochondria, their impact on cellular metabolism, and their potential for co-delivery of synergistic therapeutic agents, but these new mechanistic insights of SNs behaviour in healthy and cancer cells provide a valuable starting point. Understanding these pathways will be crucial for advancing SNs in clinical applications and maximizing their therapeutic efficacy.

## 5. Conclusions

The rapid and active penetration of cells by SNs through energy-dependent mechanisms has been demonstrated. Cytotoxicity studies indicate that SNs might have a cancer-specific cytotoxic effect on A549 lung cancer cells compared to normal cells, and that cell death occurs primarily through apoptosis at cytotoxic concentrations. Intracellularly, SNs show co-localization with lysosomes and, to a greater extent, with mitochondria. Toxicity studies at cytotoxic concentrations reveal potential damage to both organelles, with lysosomal membrane permeability associated with autophagic flux blockage and mitochondrial outer membrane permeabilization.

Moreover, RSV, selected as an antitumor agent capable of stimulating apoptosis via the mitochondrial apoptotic pathway, has been successfully encapsulated in SNs. The reduction in IC_50_ value suggests that a lower dose of SNs-RSV is needed for the same therapeutic effect as RSV alone. This implies enhanced pharmacological activity of RSV when encapsulated in SNs, possibly due to improved bioavailability, increased cellular uptake, and targeted delivery.

In summary, the co-localization of SNs with lysosomes and mitochondria, coupled with their effects on lysosomal and mitochondrial permeability, along with their potential for encapsulating mitochondria-targeted drugs, like RSV, present new avenues for targeted drug delivery and the modulation of cellular processes for positive therapeutic outcomes. These findings contribute to the advancement of nanoparticle-based drug delivery systems and hold promise for more effective treatments across various medical applications. However, further research and testing are essential to validate these benefits and ensure the safety and efficacy of the nanoparticles in diverse clinical scenarios.

## 5. Conclusions

The rapid and active penetration of cells by SNs through energy-dependent mechanisms has been demonstrated. Cytotoxicity studies indicate that SNs might have a cancer-specific cytotoxic effect on A549 lung cancer cells compared to normal cells, and that cell death occurs primarily through apoptosis at cytotoxic concentrations. Intracellularly, SNs show co-localization with lysosomes and, to a greater extent, with mitochondria. Toxicity studies at cytotoxic concentrations reveal potential damage to both organelles, with lysosomal membrane permeability associated with autophagic flux blockage and mitochondrial outer membrane permeabilization.

Moreover, RSV, selected as an antitumor agent capable of stimulating apoptosis via the mitochondrial apoptotic pathway, has been successfully encapsulated in SNs. The reduction in IC_50_ value suggests that a lower dose of SNs-RSV is needed for the same therapeutic effect as RSV alone. This implies enhanced pharmacological activity of RSV when encapsulated in SNs, possibly due to improved bioavailability, increased cellular uptake, and targeted delivery.

In summary, the co-localization of SNs with lysosomes and mitochondria, coupled with their effects on lysosomal and mitochondrial permeability, along with their potential for encapsulating mitochondria-targeted drugs, like RSV, present new avenues for targeted drug delivery and the modulation of cellular processes for positive therapeutic outcomes. These findings contribute to the advancement of nanoparticle-based drug delivery systems and hold promise for more effective treatments across various medical applications. However, further research and testing are essential to validate these benefits and ensure the safety and efficacy of the nanoparticles in diverse clinical scenarios.

## Author Contributions

Conceptualization, M.D.L.F., I.M., J.G.F.; methodology, I.M., J.G.F., M.D.L.F.; validation, E.R.D and J.G.F.; formal analysis, E.R.D.; investigation, E.R.D, I.M-J., M.I.B.; resources, M.D.L.F.; data curation, E.R.D.; writing—original draft preparation, E.R.D., J.G.F.; writing—review and editing, J.G.F., M.D.L.F.; visualization, E.R.D. and J.G.F.; supervision, I.M., J.G.F. and M.D.L.F.; project administration, M.D.L.F.; funding acquisition, M.D.L.F. All authors have read and agreed to the published version of the manuscript.

## Data Availability Statement

We encourage all authors of articles published in MDPI journals to share their research data. In this section, please provide details regarding where data supporting reported results can be found, including links to publicly archived datasets analyzed or generated during the study. Where no new data were created, or where data is unavailable due to privacy or ethical restrictions, a statement is still required. Suggested Data Availability Statements are available in section “MDPI Research Data Policies” at https://www.mdpi.com/ethics.

